# Manipulating mitochondrial dynamics in the NTS prevents diet-induced deficits in brown fat morphology and glucose uptake

**DOI:** 10.1101/2023.01.04.522581

**Authors:** Arianna Fozzato, Lauryn E. New, Joanne C. Griffiths, Bianca Patel, Susan A. Deuchars, Beatrice M. Filippi

## Abstract

Brown adipose tissue (BAT) uptakes and metabolises both glucose and triglycerides to produce heat and is activated by the central nervous system (CNS) through direct noradrenergic sympathetic innervation. Dysregulation of signalling modules in selective CNS areas such as the nucleus of tractus solitarius (NTS) are linked with altered BAT activity, obesity and diabetes. High-fat diet (HFD)-feeding increases mitochondrial fragmentation in the NTS triggering insulin resistance, hyperphagia and weight gain. Here we sought to determine whether changes in mitochondrial dynamics in the NTS can affect BAT glucose uptake. Our findings demonstrated that short-term HFD feeding reduces BAT’s ability to take up glucose, as measured by PET/CT scan. However, inhibiting mitochondrial fragmentation in NTS-astrocytes of HFD-fed rats improved BAT glucose uptake while lowering blood glucose and insulin levels. Compared with HFD-fed rats, HFD fed animals, where mitochondrial fragmentation was inhibited in the NTS-astrocytes, had higher levels of catecholaminergic innervation of BAT, and did not present HFD-dependent infiltration of enlarged white fat droplets in the BAT. In regular chow-fed rats, increasing mitochondrial fragmentation in the NTS-astrocytes reduced BAT glucose uptake, catecholaminergic innervation and β3-adrenergic receptor levels. Our data suggest that targeting mitochondrial dynamics in the NTS-astrocytes could be a beneficial strategy to increase glucose utilization and protect from developing obesity and diabetes.

## 1. Introduction

According to the World Health Organization, more than 1.9 billion people were overweight or obese worldwide in 2016 [1]. Obesity increases the likelihood of developing insulin resistance and diabetes, hence the critical need for new approaches to prevent hyperphagia and weight gain.

The brown adipose tissue (BAT) uptakes and metabolises both glucose and triglycerides to produce heat [2]. Production of heat through adaptive thermogenesis is a way for the organism to dissipate energy and maintain temperature homeostasis and burn the energy excess. [3]. In a cohort study of over 10,000 adult humans, only 7.4% were considered BAT positive, where the presence of BAT was independently associated with lower blood glucose and lower prevalence of T2DM [4]. It is estimated that, when present, 50 g of maximally stimulated BAT could produce 20% of daily energy expenditure in humans [5]. In HFD-fed rodents, BAT activity progressively decreases with the worsening of the obese phenotype [6], while in humans there is a marked decrease in BAT volume associated with obesity [7]. Genetic ablation of BAT in mice triggers obesity and increases body lipid storage [8].

BAT is densely vascularized and innervated by the sympathetic branch of the autonomic nervous system [5]. Sympathetic preganglionic neurones (SPNs) are activated by descending projections from the Raphe nuclei to SPNs, which in turn stimulate postganglionic sympathetic neurones in the sympathetic chain ganglia (SCG) that innervate BAT and release norepinephrine (NE). In rodents, this NE binds to the ꞵ3-adrenergic receptor that activates Gs-coupled G-protein coupled receptors (GPCRs) which in turn activate adenylyl cyclase, resulting in the release of cyclic AMP (cAMP) subsequently leading to triglyceride hydrolysis and release of free fatty acids. This ultimately activates uncoupling protein 1 (UCP1)-driven thermogenesis, during which heat is produced instead of adenosine triphosphate (ATP) [9].

The dorsal vagal complex (DVC) in the brainstem is an important brain area that can sense changes in hormones and nutrient levels and trigger a neuronal relay to decrease hepatic glucose production (HGP), food intake and body weight [10–14]. A key area of the DVC, the Nucleus of the tractus solitarius (NTS), plays an important regulatory role in modulating BAT activity. The NTS provides inhibitory innervation of the intermediate reticular nucleus (IRt) and the parvocellular reticular formation (IRt/PCRt), areas of the brain that innervate VGLUT3 expressing pre-sympathetic neurones in the Raphe nucleus and inhibits BAT thermogenesis induced by cooling [9,15]. Interestingly, this inhibitory action of the NTS is affected by HFD-feeding [16].

HFD-feeding causes altered mitochondrial dynamics in different brain areas and also in peripheral organs [17–21]. In the DVC, increased mitochondrial fission through activation of Dynamin-Related Protein 1 (Drp1) triggers insulin resistance [22]. Inhibition of Drp1 in the DVC restores the ability of insulin to regulate glucose metabolism and decreases body weight and food intake in HFD-fed rats, while expressing active Drp1 (Drp1-S637A) in the DVC of rats fed with regular chow causes insulin resistance, hyperphagia and body weight gain [22,23]. In our recent work, we have specifically targeted GFAP-expressing astrocytes of the NTS and expressed a dominant-negative form of Drp1 (Drp1-K38A) [22,23] which protected HFD-fed rats from developing insulin resistance, body weight gain and hyperphagia [22,23]. Indeed, astrocytes are important sensors of changes in nutritional status, with HFD-feeding they exhibit altered morphology and proliferation, which triggers astrogliosis [24], while direct chemogenetic activation of astrocytes in the NTS can cause hyperphagia and body weight gain [25].

The NTS in the DVC is important in regulating BAT activity and is highly affected by HFD-feeding and obesity. In parallel, changes in mitochondrial dynamics in the NTS astrocytes can affect insulin sensitivity and the way the NTS regulate glucose metabolism, food intake and body weight [26]. Here we investigated whether modulating mitochondrial dynamics in the NTS astrocytes can affect BAT glucose uptake and whether inhibition of mitochondrial fission in the NTS-astrocytes could improve BAT functions in HFD-fed rats.

## 2. Results

### 2.1. Short-term HFD-feeding impairs BAT glucose uptake

We first investigated whether, in our 2 week-HFD-fed model, there are any changes in BAT activity, measured as glucose uptake in dynamic fluorodeoxyglucose (^18^F) (^18^F-FDG)-PET/CT scans (Figure 1A). At thermoneutrality, control regular chow (RC)-fed rats and HFD-fed rats both expressing GFP in the NTS of the DVC (Figure 1B) presented no difference in ^18^F-FDG uptake in BAT tissue (Figure 1C), thus suggesting that basal BAT glucose uptake is not affected by HFD-feeding. As noradrenergic stimulation is the main driver of BAT thermogenesis in vivo, we tested whether the BAT response to injection of a selective ꞵ3-adrenergic agonist, CL 316243 [27] was affected by HFD-feeding. Upon injection into RC-fed rats, BAT glucose uptake increased ∼34 times, from 0.144 % of ^18^F-FDG uptake per gram of tissue in a thermoneutral unstimulated state to 4.9 % after 1mg/kg CL 316243 injection, as measured by gamma counting at the end of the scan (t: 4800s) (Figure 1C). On the other hand, HFD-fed rats did not respond to the CL 316243 injection and the percentage of ^18^F-FDG uptake per gram of tissue in BAT was similar in the thermoneutral control and CL 316243 injected rats (Figure 1C). Analysis of dynamic scans also showed that over time, there was a significant increase in the accumulation of ^18^F-FDG in the BAT of RC-fed rats treated with CL 316243 when compared with thermoneutral control RC-fed rats or HFD-fed rats that were either thermoneutral control or treated with CL 316243 (Figure 1D, E to H and Fig. S1Bi to iv).

**Figure 1:**
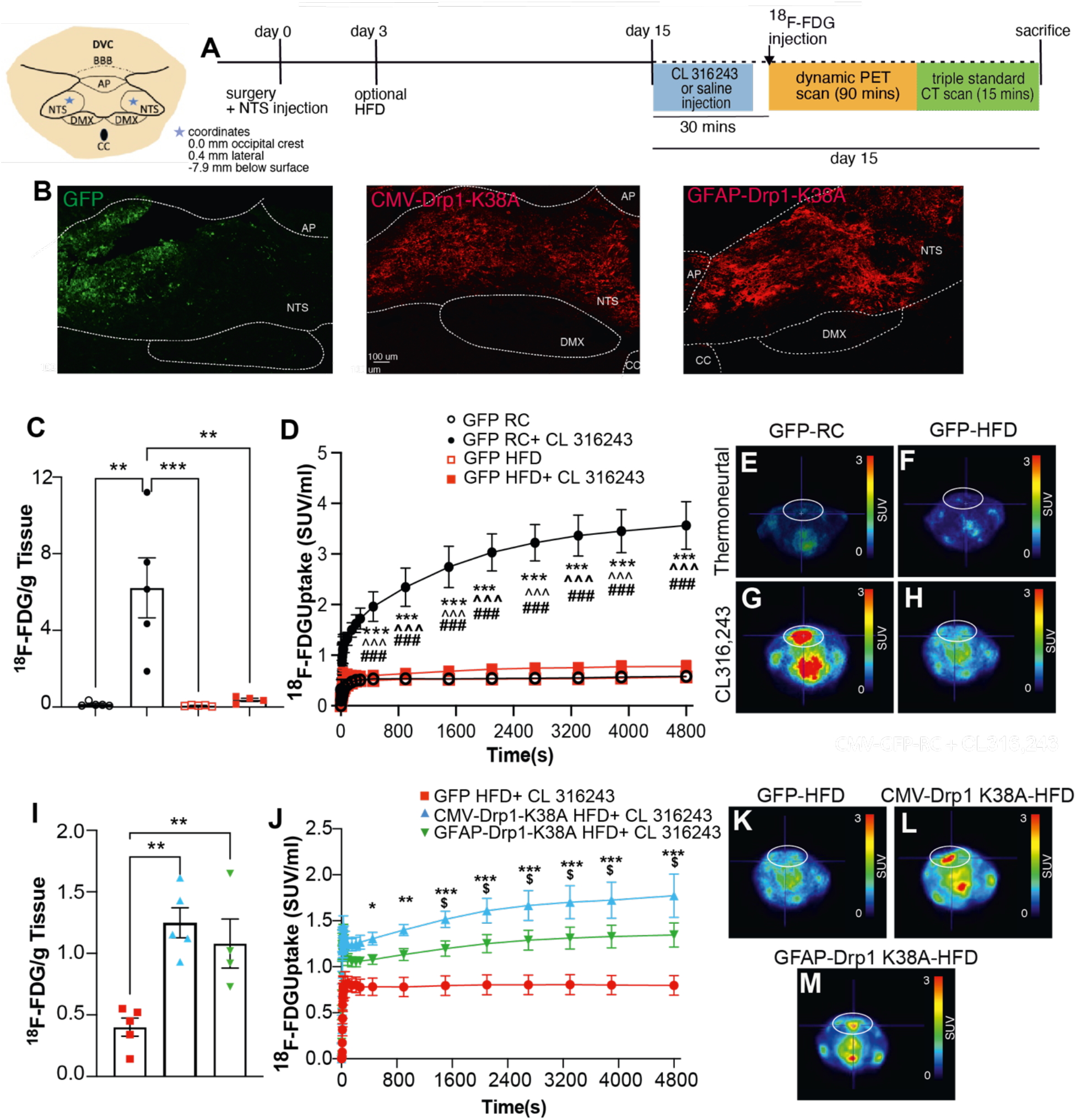
Inhibition of mitochondrial fission in the NTS prevents a reduction of dynamic glucose uptake in BAT in HFD-fed rats. (**A**) The experimental design of the PET/CT scan study. (**B**) Representative confocal images of the DVC areas expressing GFP (CMV-GFP, left) and FLAG-tagged dominant-negative Drp1 mutants CMV-Drp1-K38A (middle) and GFAP-Drp1-K38A (right) in the NTS. NTS= nucleus tractus solitarius; AP= area postrema; CC= central canal; DMX=dorsal motor nucleus of the vagus. Scale bar= 100 um. (**C**) Percentage of ^18^F-FDG uptake in BAT over the total injected corrected for ^18^F decay over time (half-life=109.7 minutes), per gram of tissue of two independent GFP RC and GFP HFD groups at thermoneutral state (n=5 for both groups mix of CMV and GFAP driven GFP expressing viruses) and following injection of 1 mg/kg CL 316243 (n=5 for both groups) as measured by gamma counting. (**D**) Graph of ^18^F-FDG accumulation in BAT over time across the experimental groups (n=3 per group). Figure 1C and D shear the same Key. (**E-H**) Representative PET scan expressed in SUV (Standard uptake value)/ml transversal images of: GFP RC animal at thermoneutrality (E), or treated with CL 316,243 (G); and of GFP HFD animals at thermoneutrality (F) or treated with CL 316243 (H). (**I)** Bar chart of the ^18^F-FDG uptake in BAT per gram of tissue as measured by gamma counting in GFP HFD (n=5), CMV-Drp1-K38A HFD (n=5) and GFAP-Drp1-K38A HFD (n=4) following injection of 1 mg/kg CL 316243 (see also supplementary Figure S1A and Bi to iv). (**J**) Graph of ^18^F-FDG accumulation in BAT over time across the experimental groups (n=3 per group). Figure 1 I and J shear the same key (**K-M**) Representative PET scan expressed in SUV/ml transversal images of: GFP HFD (K), CMV-Drp1-K38A (L) GFAP-Drp1-K38A (M) animal treated with CL 316,243 (see also supplementary Figure S1 A and Biv to vi). Circles highlight BAT area. All data were tested for normality prior to statistical tests using the Shapiro-Wilk normality test. Statistical test: Two-way ANOVA, post-hoc Tukey (C, D and J) or 1 way ANOVA, post-hoc Tukey (I). ∗p<0.05; ∗∗p<0.01; ∗∗∗p<0.001. Where applicable values are shown as mean ± SEM and single data point highlighted. In D and J Statistical significance is shown from 270s to 4,800. In D, * p value for GFP RC vs GFP RC+ CL 316,243; ^ p value for GFP RC+ CL 316243 vs GFP HFD; # p value for GFP RC+ CL 316,243 vs GFP HFD+ CL 316,243. In J, * p value for GFP RC+ CL 316243 vs CMV-Drp1-K38A HFD+ CL 316,243; $ p value for GFP HFD+ CL 316243 vs GFAP-Drp1-K38A HFD + CL 316243.

### 2.2. Inhibition of mitochondrial fission in the NTS improves BAT glucose uptake in HFD-fed rats

Short-term HFD-feeding increases mitochondrial fission in the DVC thus triggering insulin resistance, hyperphagia and body weight gain [22,23]. Inhibition of mitochondrial fission in the NTS by expressing a dominant negative Drp1-K38A via adenoviral delivery prevented HFD-dependent insulin resistance and decreased body weight and food intake [22,23]. We tested whether a further protective effect is associated with changes in BAT glucose uptake. Rats were injected with adenoviruses expressing Drp1-K38A under either the ubiquitous CMV promoter to target any cell in the NTS or under the GFAP promoter to target GFAP-positive astrocytes only (Figure 1B) [22,23]. Rats were kept for 2 weeks with HFD and then injected with CL 316,243 and ^18^F-FDG to undergo PET/CT scan analysis (Figure 1A). Inhibition of mitochondrial fission in the NTS of the DVC in HFD-fed rats was sufficient to promote higher levels of ^18^F-FDG uptake in BAT compared to the HFD-fed control group as shown by the different levels of ^18^F-FDG accumulation at the end of the scan (Figure 1I). This effect was similar when Drp1-K38A was expressed non-specifically in both neurones and glial cells of the NTS and when Drp1-K38A was expressed only in GFAP-expressing astrocytes. Analysis of dynamic scans also showed that over time, there were significantly higher levels of accumulation of ^18^F-FDG in the BAT of both the CMV- and GFAP-Drp1-K38A expressing models compared to HFD-fed control group (Figure 1J, K to M and Fig. S1Biv to vi), thus suggesting a partial restoration of BAT ability to uptake glucose in these animals compared to HFD controls.

### 2.3. Effect of diet and mitochondrial dynamics on white adipose tissue glucose uptake and accumulation

Improvement in the metabolic health can be also associated with increased WAT glucose uptake [28]. At thermoneutrality there was no significant difference between the ^18^F-FDG uptake in WAT of RC-fed and HFD-fed animals (Figure 2A), while upon CL 316243 stimulation, there was a significant increase in ^18^F-FDG uptake in WAT of regular chow but not in HFD-fed animals, as measured by gamma counting (Figure 2A). Interestingly, inhibition of mitochondrial fission in the DVC of HFD-fed rats did not increase ^18^F-FDG uptake in WAT in either CMV-Drp1-K38A or GFAP-Drp1-K38A models (Figure 2B).

**Figure 2:**
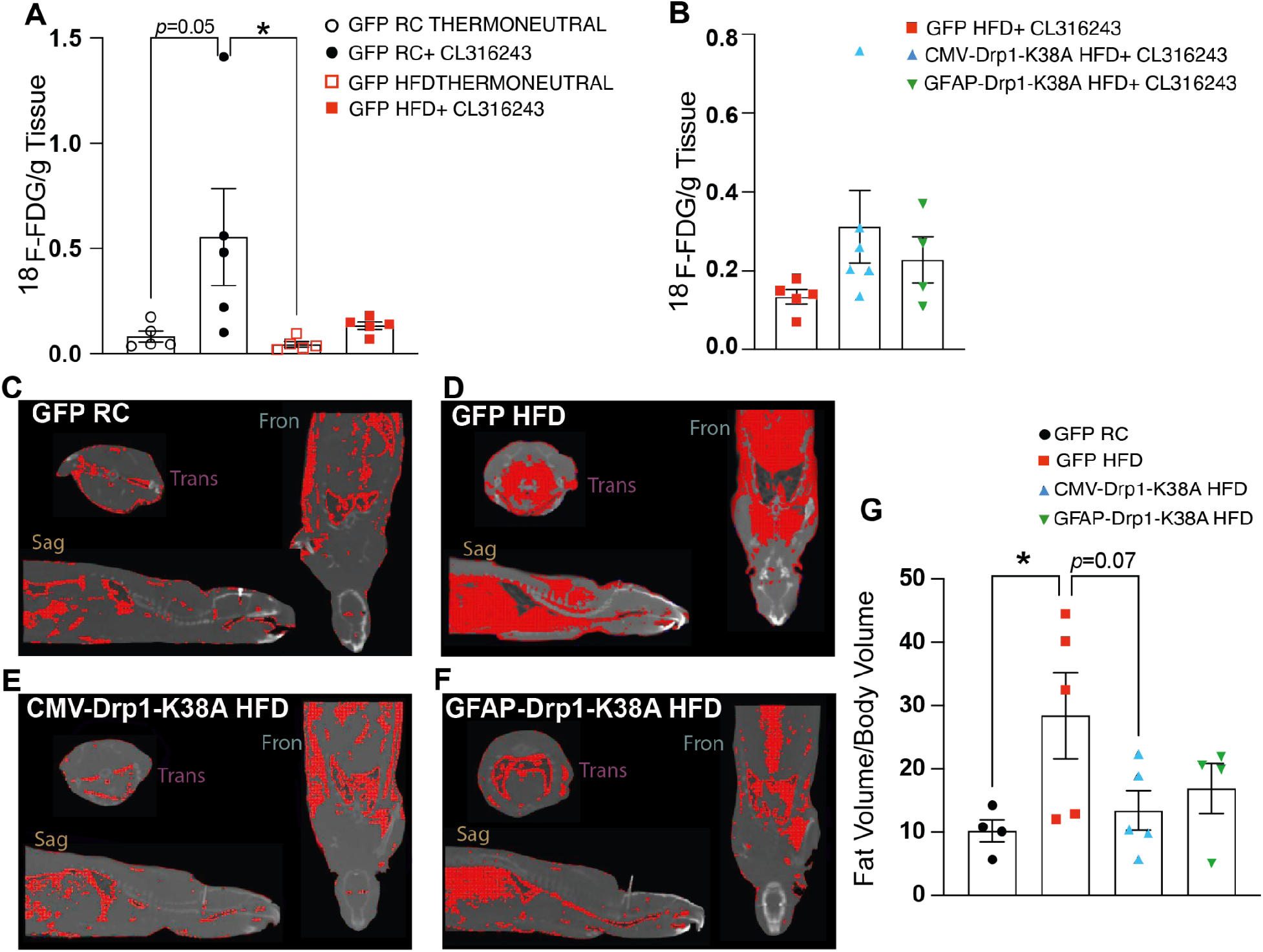
Inhibition of mitochondrial fission in the NTS of HFD-fed rats does not affect the volume of WAT or WAT dynamic glucose uptake. (**A-B**) Percentage of injected dose of ^18^F-FDG in WAT per gram of tissue as measured by gamma counting. (A) RC vs HFD-fed groups at thermoneutrality without stimulation or injected with 1 mg/kg CL 316243 (n=5 for CMV-GFP RC, CMV-GFP HFD). (B) CMV-GFP HFD, CMV-Drp1-K38A HFD (n=6) and GFAP-Drp1-K38A HFD (n=4) injected with 1 mg/kg CL 316243. (**C-F**) Representative composite image CT scan (greyscale) and adipose tissue volume (red) of (C) GFP RC, (D) GFP HFD, (E) CMV-Drp1-K38A HFD and (F) GFAP-Drp1-K38A HFD. (**G**) Adipose tissue/total volume ratio for all groups (n=5 for GFP RC, GFP HFD and CMV-Drp1-K38A HFD, and n=4 for GFAP-Drp1-K38A HFD). All data were tested for normality prior to statistical tests using the Shapiro-Wilk normality test. Statistical test: 2-way ANOVA with Tukey post-hoc (A) or One-way ANOVA with Dunnet post-hoc (B and G). Values are shown as mean ± SEM and single data points are highlighted. ∗p < 0.05.

Information obtained with the CT scans also allowed quantification of fat volume. We calculated the ratio between total body volume and adipose tissue volume by exploiting the differences in the density of tissues expressed in Hounsfield Units. RC-fed animals presented a low level of visceral and subcutaneous adiposity when compared with HFD-fed rats, suggesting that 2-weeks of HFD are sufficient to induce profound changes in visceral and subcutaneous adiposity in adult male rats (Figure 2C-G). Inhibition of mitochondrial fission in neurones and glial cells of the NTS of HFD-fed rats caused a trend towards a decrease in visceral and subcutaneous adiposity when compared to HFD-fed control but this did not reach significance; the same trend was observed when only astrocytes were targeted (Figure 2C-G).

### 2.4. Inhibition of mitochondrial fission in NTS astrocytes prevents WAT-like droplet infiltration into BAT and increases TH-positive inputs onto BAT

Our PET scan data clearly suggest that blocking mitochondrial fission in astrocytes is sufficient to mitigate an HFD-dependent decrease in BAT glucose uptake seen in control HFD-fed animals (Figure 1). To understand the mechanisms behind these observations, we analysed BAT weight and morphology. Looking at BAT weight in our PET scan animals there were no differences between the groups (Fig. S1C). For the morphological analysis, since the PET animals were treated with the agonist, in order to avoid confounding factors in our analysis, we used samples from HFD-fed rats expressing either GFAP-GFP or GFAP-Drp1-K38A in the NTS generated by Patel et (2021). Drp1-K38A-expressing HFD-fed rats showed a decrease in body weight, food intake and fat accumulation when compared with GFP-expressing HFD-fed rats [23] and had similar BAT weights (Figure 3A). However, H&E analysis showed high levels of enlarged lipid droplets in the BAT of HFD-fed rats (Figure 3B). While BAT usually exhibits small multilocular droplets in a single cell, the presence of big white droplets could suggest whitening and hypertrophy of the BAT. Expression of Drp1-K38A in astrocytes prevented large fat droplet accumulation and exhibited a higher number of small droplets per field (Figure 3B).

**Figure 3:**
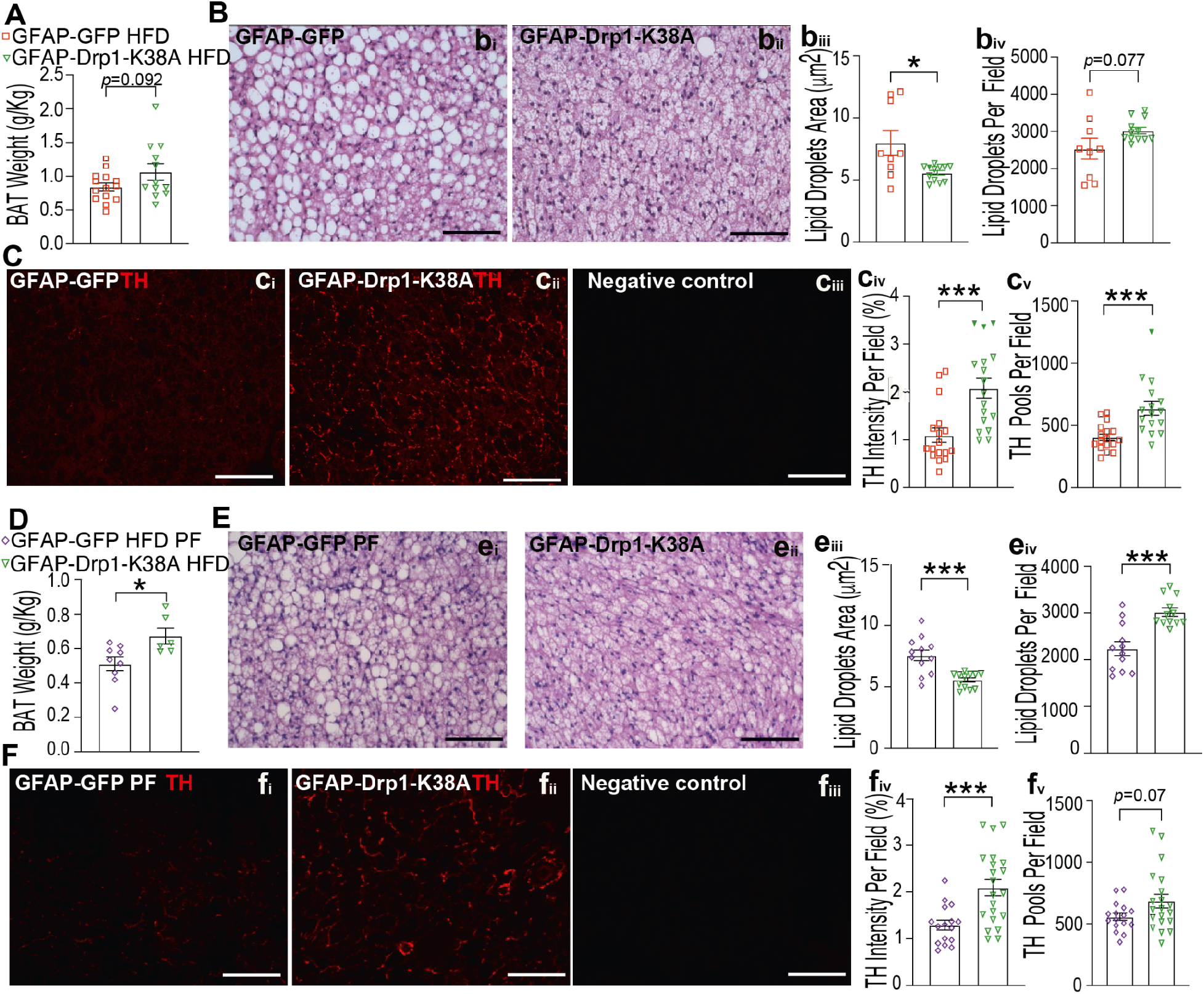
Inhibition of mitochondrial fission in the NTS-astrocytes of HFD-fed rats prevents hypertrophy and increases noradrenergic innervation of BAT. (**A and D**) BAT weight normalised to body weight (g/kg) in GFAP-GFP (n=14) and GFAP-Drp1-K38A (n=12) animals fed with HFD (cohorts from Patel et al., 2021) (A) and GFAP-GFP pair-fed (PF) (n=9) and GFAP-Drp1-K38A (n=6) HFD-fed animals (D). (**B and E**) Representative images of H&E staining of serial BAT sections cut at 5 um for GFAP-GFP HFD (bi) and GFAP-Drp1-K38A HFD (bii) animals and for GFAP-GFP PF (ei) and GFAP-Drp1-K38A HFD (eii) animals. (biii and eiii) Lipid droplet area quantification for the indicated groups and (biv and eiv) lipid droplets number per visual field for the indicated groups. GFAP-GFP HFD (n=3, 3 technical repeats), GFAP-Drp1-K38A HFD (n=4,3 technical repeats), GFAP-GFP HFD PF (n=3, 3 technical repeats.) and GFAP-Drp1-K38A-PF HFD (n=4, 3 technical repeats). (**C and F**) Representative confocal images of tyrosine hydroxylase (TH) staining in serial BAT sections cut at 5um for GFAP-GFP HFD group (ci) and GFAP-Drp1-K38A HFD group (cii) and for GFAP-GFP PF (fi) and GFAP-Drp1-K38A HFD group (fii). (ciii and fiii) Negative control (- primary antibody). (civ and fiv) TH intensity quantification per visual field for the indicated groups and (cv and fv) number of TH pools per visual field for the indicated groups. GFAP-GFP animals (n=4, 4 technical repeats) and GFAP-Drp1-K38A animals (n=4, 4 technical repeats), GFAP-GFP HFD PF animals (n=4, 4 technical repeats) and GFAP-Drp1-K38A animals (n=4, 4 technical repeats). Statistical test: unpaired, 2 tails T-test. Values are shown as mean ± SEM and single data points are highlighted. ∗p < 0.05; ∗∗∗p < 0.001.

Changes in BAT morphology and in fat droplet accumulation could be correlated with altered BAT activity. Activation of the sympathetic inputs onto BAT and thus ꞵ-adrenergic receptor stimulation is the most potent trigger for BAT activation to promote thermogenesis [29]. In order to determine whether alterations of the sympathetic innervation of BAT are associated with decreased BAT glucose uptake in HFD-fed rats, we examined staining for tyrosine hydroxylase (TH) which labels catecholaminergic neurones. Neuronal projections that terminate in the BAT could therefore be visualised with an anti-TH antibody and the intensity and number of TH+ pools quantified. We observed low levels of TH staining with HFD-feeding (Figure 3C) suggesting altered neuronal projections to the BAT. Interestingly, inhibiting mitochondrial fission in the NTS astrocytes was sufficient to cause higher TH levels in BAT compared to control HFD-fed rats, as suggested by an increase in TH intensity and in the number of pools within BAT (Figure 3Ci, Civ-v).

Our data so far suggest that inhibition of mitochondrial fission in the NTS astrocytes of HFD-fed animals prevents formation of WAT-like lipid droplet and increases catecholaminergic terminals availability in the BAT. Compared with the HFD-fed GFP rats, the Drp1-K38A HFD-fed rats also presented significantly lower body weight, food intake and fat accumulation [23]. However, whether the protective effect on BAT was due to a decrease in body weight or due to altered NTS signalling is not clear. To answer this question, we performed a pair-feeding study where HFD-fed rats expressing GFP (GFP-PF) in the NTS astrocytes were fed with an average amount of food that rats expressing Drp1-K38A in the NTS astrocytes (Drp1-K38A) normally eat daily (Fig. S2A). Cumulative food intake and body weight were similar between the two cohorts (Fig. S2B and C). Interestingly, there was a significantly higher body fat accumulation in GFP-PF rats compared with the Drp1-K38A ones (Fig. S2D) while the BAT volume was lower (Figure 3D). H&E data showed a larger fat droplet size in GFP-PF rats and a lower numbers of lipid droplets (Figure 3E), thus demonstrating this change in droplet size and number occurs independently of food intake and body weight gain, which could be suggestive of a centrally-driven mechanism.

The TH analysis showed that GFP-PF rats have lower TH staining intensity and TH pool number compared with the Drp1-K38A HFD rats (Figure 3F), thus suggesting that the protective effect that inhibition of mitochondrial fission in NTS astrocytes has on the ꞵ-adrenergic sympathetic pathway is independent of diet and body weight.

### 2.5. Fatty acid metabolism and inflammatory markers in the BAT can be modulated by alterations in mitochondrial dynamics in NTS astrocytes

To understand if changes in NTS mitochondrial dynamics can affect peripheral BAT metabolism and inflammation, we analysed the expression levels of multiple genes involved in different aspects of BAT metabolism and inflammation (Figure 4). When looking at fatty acid oxidation and lipolysis, compared with rats expressing GFP, rats expressing Drp1-K38A specifically in NTS astrocytes showed lower levels of lipolytic enzymes AGTL (*Pnpla2*) and HSL (Adipose Triglyceride Lipase and Hormone-Sensitive Lipase, hsl) [30] and higher levels of *Cd36*, a scavenger receptor that favours the uptake of long chain fatty acids and exogenous Coenzyme Q (CoQ)[2,31]. There were also significantly lower levels of the ER-stress marker C/EBP homologous protein, CHOP (*Dditn3*) [32], as well as inflammatory markers *NFK-*ꞵ *(NfkB)* and *TNF-*⍺ (*TnfA*) in Drp1-K38A HFD rats (Figure 4 A). Interestingly, genes associated with insulin and glucose metabolism were unchanged. Among the markers of BAT activation, gene encoding for cell death-inducing DFFA Like Effector A (*Cidea)[33]* was lower while ꞵ3-adrenergic receptor (*Adrb3*) was moderately higher but this did not reach significance. *Ucp1*, PGC1-⍺ (*Pprgc1a)*) and PPAR ɣ (*Pprg* ɣ) [29] were not altered in normothermic conditions (Figure 4A). Injection of CL 316243 to activate BAT glucose uptake triggered an increase in PGC1-⍺ in both GFP and Drp1-K38A HFD rats while only the latter group presented increased UCP1 levels upon stimulation. Interestingly, most of the changes in gene expression associated with the Drp1-K38A HFD group were lost in the GFP pair-fed group (Figure 4C), thus suggesting changes in BAT gene expression occur in a feeding-dependent manner. Of note, *Cidea* levels were lower only in Drp1-K38A animals while PF and HFD control presented higher levels, thus suggesting that this gene is regulated independently of feeding and body weight. Interestingly both PGC1-⍺ and insulin receptors (*Insr)* were elevated in PF animals only (Figure 4C). Collectively this analysis suggests that by altering mitochondrial dynamics in the NTS we can change the metabolic status of the BAT thus altering its ability to metabolise glucose and lipids and produce heat.

**Figure 4:**
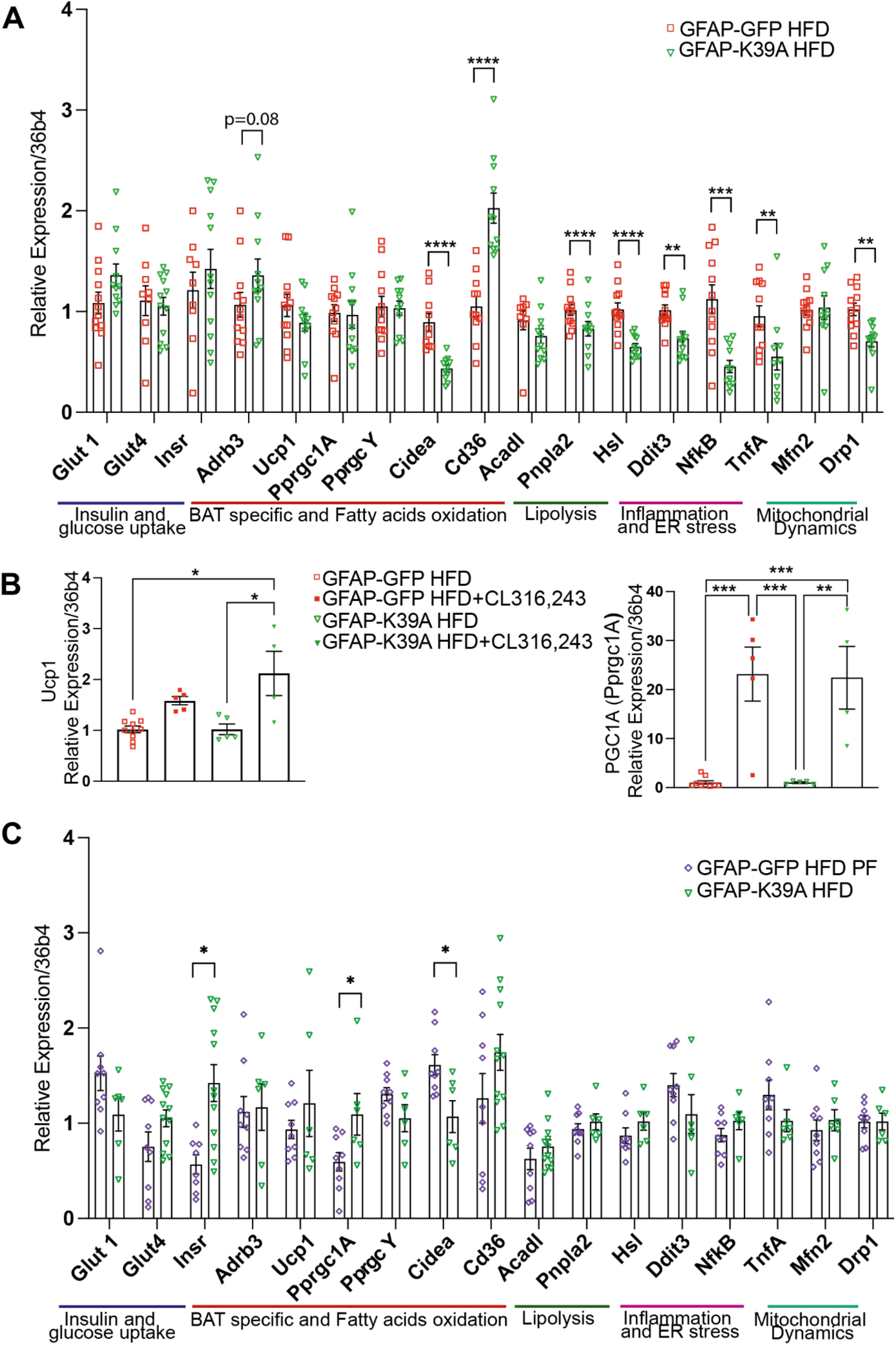
Changes in BAT gene expression associated with altered mitochondrial dynamics in the NTS. qPCR on BAT tissue against five classes of genes related to insulin and glucose metabolism, BAT specific and fatty acid oxidation, lipolysis, inflammation and ER stress and mitochondrial dynamics. Indicated gene expression levels are shown relative to the expression of the housekeeping gene 36b4. (**A**) Relative expression of genes of interest in GFP HFD (n=11) versus Drp1-K38A HFD (n=11) animals (samples from Patel et al 2021). (**B**) Relative expression of genes involved in thermogenesis in HFD-fed control GFAP-GFP (n=10) and GFAP-Drp1-K38A (n=5) injected or not with CL 316243. **(C)** Relative expression of genes of interest in GFP-HFD-PF (n=9) versus Drp1-K38A HFD (n=6) animals. Cohorts were run at separate times with internal controls to ensure gene expression consistency. Statistical test: unpaired, two tails T-test in A and C and two-way ANOVA with Tukey post hoc in B. Values are shown as mean ± SEM and single data points are highlighted. ∗p < 0.05; ∗∗p < 0.01; ∗∗∗p < 0.001; ∗∗∗∗p < 0.0001.

### 2.6. Increased mitochondrial fission in NTS astrocytes causes lower BAT activity compared to control

We recently showed that increasing mitochondrial fission in RC-fed rats, by expressing a constitutively active form of Drp1 (Drp1-S637A) under the CMV promoter in the NTS (Figure 5A), results in hyperphagia, higher weight gain and greater abdominal fat accumulation compared to RC control rats expressing GFP in the NTS [23]. We first determined whether the expression of Drp1-S637A only in astrocytes of the NTS was sufficient to cause a similar phenotype. To this aim, we developed and injected an adenovirus expressing Drp1-S637A under the GFAP promoter in the NTS of RC-fed rats using GFP as control (Figure 5B). After 2 weeks, Drp1-S637A rats presented greater cumulative food intake and body weight (Fig. S3A and B), while no changes in fat accumulation was seen (Fig. S3C). Interestingly insulin levels were higher in the Drp1-S637A cohort (Fig. S3D) while expression of Drp1-K38A in the NTS astrocytes of HFD-fed rats caused lower insulin levels (Fig. S2E). Moreover, blood glucose levels were significantly lower in Drp1-K38A animals when compared to both GFP-HFD and GFP-PF-HFD groups (Fig. S2F), whilst they were unchanged in Drp1-S637A group compared to GFP-RC controls (Fig. S3E). Collectively these data suggest that is sufficient to increase mitochondrial fission in the NTS-astrocytes to alter body weight, food intake and insulin levels.

**Figure 5:**
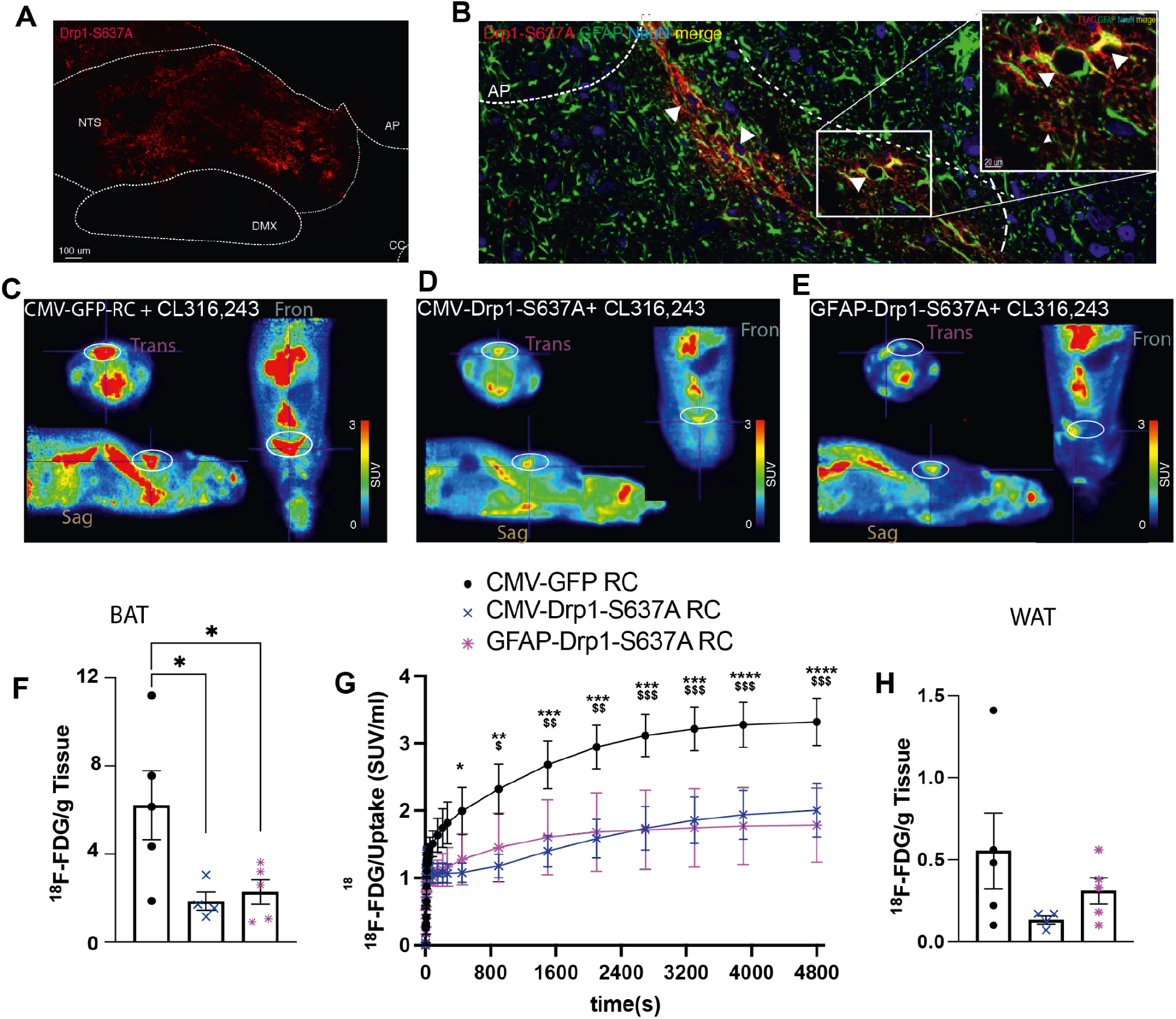
Activation of mitochondrial fission in the NTS of RC-fed rats decreases dynamic glucose uptake in BAT. (**A**) Representative confocal image of viral delivery of FLAG-tagged CMV-Drp1-S637A in the NTS of a RC-fed animal. (**B**) Representative confocal image of viral delivery of FLAG-tagged GFAP-Drp1-S637A in the NTS of a RC-fed animal. Viral expression was strictly observed in GFAP+ astrocytes (white arrows). (**C-E**) Representative PET scan expressed in SUV/ml of a CMV-GFP RC animal (C), CMV-Drp1-S637A animal (D) and GFAP-Drp1-S637A (E) animal. BAT circled in white. (**F**) Percentage of the injected dose of ^18^F-FDG in BAT per gram of tissue of CMV-GFP RC (n=5), CMV-Drp1-S637A (n=4) and GFAP-Drp1-S637A (n=5) following injection of 1 mg/kg CL 316,243 as measured by gamma counting. (**G**) ^18^F-FDG accumulation in BAT over time across the experimental groups (n=4). Statistical significance is shown from t=270s to T4800s,*=CMV-GFP RC vs CMV-Drp1-S637A); $=CMV-GFP RC vs GFAP-Drp1-K38A; (**H)** Percentage of the injected dose of ^18^F-FDG in WAT per gram of tissue of CMV-GFP RC (n=5 per group), CMV-Drp1-S637A (n=4) and GFAP-Drp1-S637A (n=5) following injection of 1 mg/kg CL 316243 as measured by gamma counting. All data were tested for normality prior to statistical tests using the Shapiro-Wilk normality test. Statistical test: One-way ANOVA, post-hoc Tukey (F, H) or Two-way ANOVA, post-hoc Tukey (G). *p < 0.05, **p<0.01, ***p=0.001, ****p=0.0001. Where applicable values are shown as mean + SEM and single data points are highlighted.

We next performed ^18^F-FDG-PET/CT scans to determine whether Drp1-S637A rats have altered BAT glucose uptake. Scans were done as described in Figure1A. Before the scans, rats were injected with 1mg/kg CL 316243 to stimulate BAT activity. Compared to RC-fed rats expressing GFP in the NTS, rats expressing Drp1-S637A (Figure 5B) exhibited lower levels of ^18^F-FDG accumulation at the end of the PET scan (t=4800s) (Figure 5C, D and F). Astrocyte-specific expression of Drp1-S637A was also sufficient to cause lower levels of BAT glucose uptake (Figure 5E and F). Analysis of the dynamic scan showed that over time, there was a significantly higher accumulation of ^18^F-FDG in the BAT of RC-fed rats when compared with both the CMV and GFAP-Drp1-S637A expressing models (Figure 5G). While we could observe a trend showing decreased ^18^F-FDG uptake in WAT of both the Drp1-S637A cohorts when compared with RC control rats, this effect was not significant due to high sample variability (Figure 5H).

### 2.7. Mitochondrial fission in NTS astrocytes decreases BAT mass and catecholaminergic innervation of BAT

We then analysed whether increasing mitochondrial fragmentation in the NTS could also affect BAT morphology and catecholaminergic projections to the BAT (Figure 6). RC-fed rats expressing Drp1-S637A in NTS astrocytes presented with lower BAT weight (Figure 6A). H&E staining analyses suggested no differences in the number and area of the lipid droplets between the GFP control and the Drp1-S637A model (Figure 6B). TH staining and the number of TH+ pools in the BAT of rats expressing Drp1-S637A in NTS-astrocytes were significantly lower when compared with the RC-GFP group (Figure 6C). These data suggest that increasing mitochondrial fission in the NTS astrocytes affects the catecholaminergic innervation of BAT, which could also reduce the ability of BAT to uptake glucose. Gene expression analysis in BAT tissues indicated lower *Adrb3*, *Ucp1* and *Cidea* levels, in line with lower BAT activity in the Drp1-S367A cohort. In addition, whilst the level of Cd36 was higher in the Drp1-K38A cohort, in the Drp1-S637A cohort this was lower (Figure 6D) indicating altered fatty acids and CoQ uptake in these animals. *TNF-⍺* was also elevated, thus suggesting higher levels of inflammation in the BAT of the Drp1-S637A expressing animals. Interestingly we could see elevated insulin receptor levels similar to the PF cohort and lower levels of Drp1, similar to the Drp1-K38A cohort (Figure 6D).

**Figure 6:**
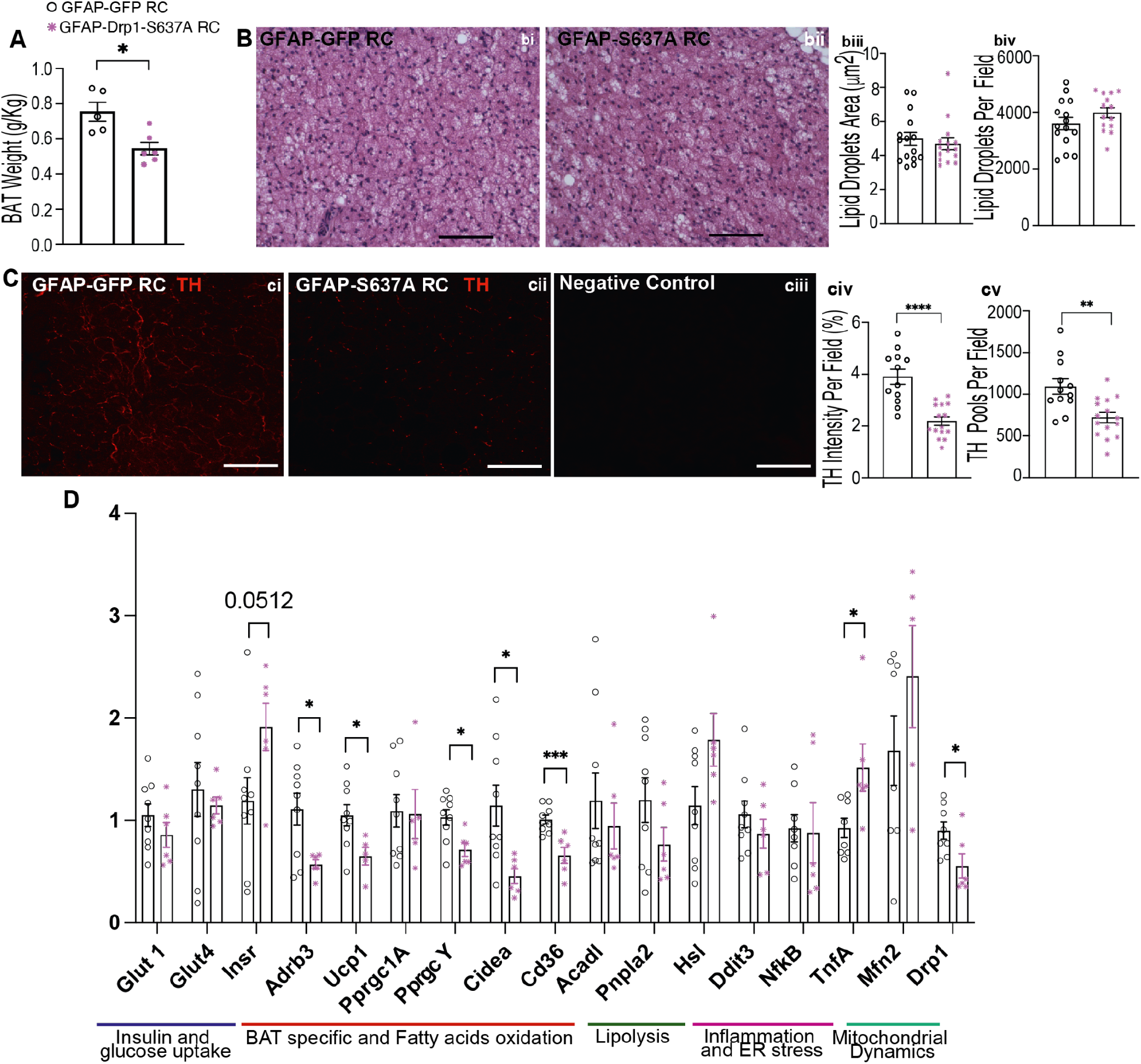
Promoting mitochondrial fission in the NTS leads to lower BAT mass and noradrenergic innervation of BAT and alters key BAT genes in regular chow-fed animals. (**A**) BAT weight normalised to body weight (g/kg) in GFAP-GFP (n=5) and GFAP-Drp1-S637A (n=6) animals fed with RC. (**B**) Representative H&E staining on serial BAT sections cut at 5 um for (bi) GFAP-GFP HFD and (bii) GFAP-Drp1-S637A. (biii) lipid droplet area quantification and (biv) lipid droplet number per visual field for GFAP-GFP HFD (n=3, 3-4 technical repeats) and GFAP-Drp1-S37A HFD (n=4, 3-4 technical repeats). (**C**) Representative confocal images of tyrosine hydroxylase (TH) staining on serial BAT sections cut at 5um for (ci) GFAP-GFP group and (cii) GFAP-Drp1-S637A group. (ciii) A representative negative control (- primary antibody). (civ) TH intensity quantification per visual field and (cv) number of TH pools per visual field for GFAP-GFP animals (n=4, 3-4 technical repeats) and GFAP-Drp1-S637A animals (n=4, 3-4 technical repeats). (**D**) Relative expression of genes of interest in GFAP-GFP (n=8) versus GFAP-Drp1-S637A (n=6) animals fed ad libitum. Statistical test: unpaired, 2 tails T-test. Values are shown as mean + SEM and single data points highlighted. ∗p < 0.05; ∗∗p < 0.01; ∗∗∗p < 0.001; ∗∗∗∗p < 0.0001.

Fatty acid transporter *CD36* is significantly upregulated in animals expressing Drp1-K38A and significantly lower in animals expressing Drp1-S637A when compared to their matching controls. Previous literature has shown that CD36 is involved in the intracellular transport of exogenous CoQ [31], a critical component of the electron transport chain whose deficiency was associated with insulin resistance in other tissues, such as the WAT and muscle in mice [34]. CoQ uptake in BAT of CD36-deficient mice was also greatly impaired [31]. Here we investigated whether the alterations in *CD36* mRNA transcripts in BAT from K38A and S637A animals are associated with changes in levels of CoQ in this tissue. Our results indicated no significant changes in CoQ levels across our experimental groups, suggesting that this mechanism is not affected in our models (Fig. S4). This analysis was carried out in tissues taken from rats kept at room temperature. It would be interesting to test whether β-adrenergic stimulation of BAT may reveal some changes in CoQ levels between the groups.

## 3. Discussion

Increasing BAT activity has the potential to increase energy expenditure, reduce body weight and improve whole-body metabolism. In this study, we showed that inhibition of mitochondrial fission in NTS astrocytes of HFD-fed rats increases BAT glucose uptake and prevents alteration of ꞵ-adrenergic stimulation while also lowering *NFK-*ꞵ and *TNF-*⍺ expression and decreasing blood glucose and insulin levels. Interestingly, increasing mitochondrial fission in NTS-astrocytes of RC-fed rats, leads to lower BAT glucose uptake, decreases ꞵ-adrenergic signalling and increases *TNF-*⍺ expression in the BAT (Figure 7). Together, our data indicate that HFD-feeding and altered mitochondrial dynamics in the NTS affect BAT regulation and could alter energy expenditure thus affecting whole-body glucose homeostasis.

**Figure 7:**
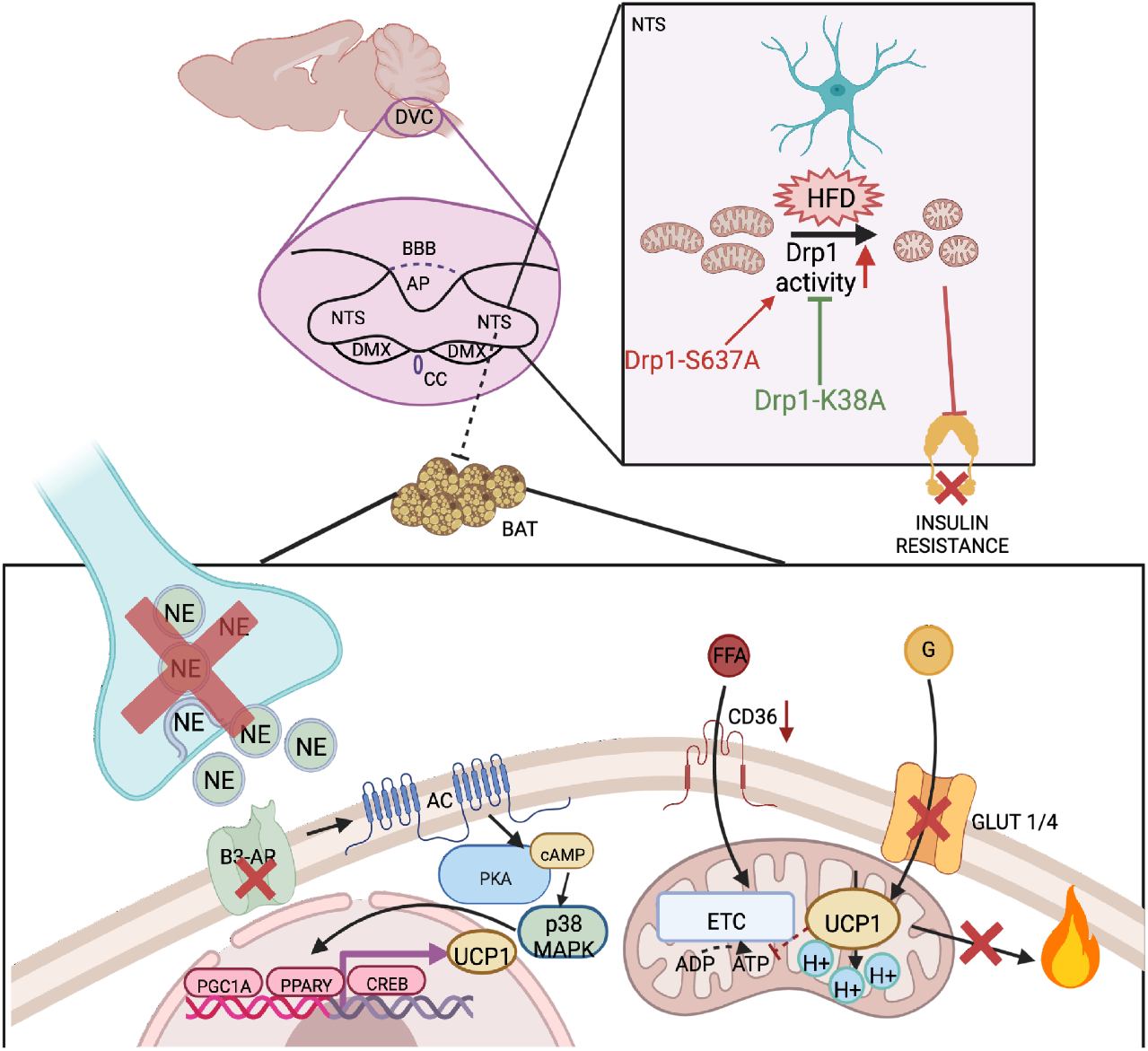
Schematic representation of working hypothesis: Short-term HFD-feeding decreases BAT ability to uptake glucose, and alters BAT morphology and neural innervation. Inhibition of mitochondrial fragmentation in the NTS-astrocytes of HFD-fed rats can prevent the effects on BAT morphology and innervation, and increase BAT glucose uptake. In regular chow-fed rats, increasing mitochondrial fragmentation in the NTS-astrocytes reduces BAT glucose uptake, alters BAT innervation, and β3-adrenergic receptor levels. Targeting mitochondrial dynamics in the NTS-astrocytes has the potential to increase glucose utilization and protect from developing obesity and diabetes.

The NTS plays an important role in modulating BAT activity. For example, vagal afferent activation inhibited the normal thermogenic responses to cooling in rats, an effect that was prevented by injections of either glutamatergic antagonists in the NTS or GABA_A_ antagonists in the rostral raphe pallidus (rRPa) [35]. Furthermore, disinhibition of neurones in the intermediate NTS ablates cooling- or febrile-evoked BAT sympathetic activation and thermogenesis [9,15,35]. Together this suggests that the NTS is activated by vagal afferent inputs and provides GABAergic inhibition of sympathetic premotor neurones in the rRPa. Importantly, in a 60-day HFD-fed obese rat model, the normal cooling-evoked increases in BAT activation are reduced and this effect was reversed by injection of glutamatergic antagonists into the NTS, suggesting that the inhibitory action of the NTS is affected by HFD-feeding [16]. Our work shows that HFD-feeding and increased mitochondrial fragmentation of the NTS-astrocytes can reduce BAT glucose uptake and decrease TH staining. Interestingly, increasing mitochondrial fission in the astrocytes of the NTS also decreased BAT mRNA levels of β3-adrenergic receptor. As ꞵ-adrenergic receptor stimulation is key to modulating BAT activity, altered noradrenergic signalling in BAT could decrease glucose uptake. A decrease in TH staining associated with an increased size of the fat droplets in the BAT of HFD-fed rats suggests potential degeneration of the BAT, and inhibition of mitochondrial fragmentation in the NTS could have a protective effect.

Astrocytes are important for sensing metabolic changes [36]. In the DVC, they can proliferate and become reactive in response to HFD-feeding and can directly alter feeding behaviour when chemo-genetically activated [25]. In the hypothalamus, short-term HFD-feeding altered astrocyte number, density, and activity [35] and ablation of insulin receptors in astrocytes affected glial morphology and mitochondrial function leading to POMC neuronal activation [37]. Here we showed that altered mitochondrial dynamics in astrocytes of the NTS can affect BAT glucose uptake and morphology. These data, and our findings, therefore suggest that astrocytes are important to maintaining metabolic health. Indeed, mice with astrocyte-specific insulin receptor knockdown (IrKO GFAP) exhibited sex-specific alteration of metabolic functions, while BAT exhibited higher adipocyte size in both sexes. Interestingly, the BAT of these animals had reduced adrenergic innervation, and lower ꞵ3-adrenergic receptor expression levels were also observed in IrKO GFAP [38]. These observations are in line with our data where alteration of mitochondrial dynamics in NTS-astrocytes can influence brown adipocyte size and ꞵ-adrenergic innervation while also affecting insulin sensitivity [23].

Astrocytes are also involved in the regulation of glutamate homeostasis [39,40,41] and HFD-feeding and obesity impair this mechanism, leading to increased brain glutamate levels, dysfunction in extra-synaptic NMDA levels and mitochondrial dysregulation [42]. Glutamate was shown to induce iNOS in the brain [43] while obesity and HFD were associated with increased iNOS levels mainly in astrocytes. Astrocytes express iNOS and inhibiting mitochondrial fission in the NTS decreased the expression of iNOS in this brain region [23]. Our study suggests a potential mechanism that may explain the link between mitochondrial dynamics in NTS astrocytes and BAT activation, where increased mitochondrial fission in the NTS can lead to higher inflammation via increased iNOS expression in astrocytes. Dysfunctional astrocytic mitochondrial dynamics may also lead to a reduced capacity of astrocyte glutamate uptake and its clearance from the extracellular space, thereby causing alterations in neurotransmission. This could eventually lead to alterations in the sympathetic innervation to BAT and consequent activation and thermogenesis. However, further investigations are required to establish a connection between astrocytes’ regulation of glutamate availability at the site of vagal afferents within the NTS, and BAT activation.

Decreasing adiposity is associated with better metabolic outcomes both in humans and rodents [44–46]. Our CT data revealed that following two weeks of HFD-feeding, rats accumulate significantly more WAT (defined as a combination of subcutaneous and abdominal depots) than RC rats. The presence of increased WAT accumulation following HFD diet was partially prevented by inhibiting mitochondrial fission in the NTS, thus suggesting that the NTS and mitochondrial dynamics play a key role in modulating fat accumulation.

Insulin receptor levels were increased in the BAT of RC-fed rats expressing Drp1-S637A in the NTS. We also found that, similarly to HFD-fed controls, Drp1-S637A-expressing animals fed with RC diet, were hyperinsulinemic. HFD causes peripheral insulin resistance, and activation of mitochondrial fission in the NTS of these animals appears to partially mimic the detrimental effects of HFD on insulin sensitivity. Thus, elevated insulin receptor gene expression in BAT could be a compensatory mechanism employed to respond to insulin resistance induced by the activation of mitochondrial fission in astrocytes within the NTS. Importantly, the inhibition of mitochondrial fission in the astrocytes of the NTS of rats prevented HFD-induced hyperinsulinemia and hyperglycaemia, and this effect was independent of food intake; in fact, HFD-paired-fed animals, similarly to *ad-libitum* HFD-fed controls, developed hyperinsulinemia and did not present a significant decrease in blood glucose. This suggests that hyperinsulinemia is not directly induced by alterations in feeding behaviour, but rather, by a centrally driven mechanism dependent upon mitochondrial dynamics within astrocytes of the NTS.

Interestingly, BAT insulin receptor levels were also elevated in the pair-fed model. While informative, the pair-feeding paradigm has limitations. One of these is the fact that the model is dependent on the ability of the animals to distribute the available food between one refeeding window and the following one, as animals may consume all their available food before being provided with more. This could therefore lead to a pattern of intermittent fasting which could affect insulin receptor gene expression in BAT, as this organ potently responds to nutritional states [47].

Altered ꞵ-adrenergic stimulation in BAT should increase *UCP1* mRNA levels. Interestingly, UCP1 levels were unchanged in thermoneutral animals fed with HFD when compared with animals treated with the ꞵ-adrenergic agonist, while a clear increase was observed in CL 316243 treated animals expressing Drp1-K38A in astrocytes of the NTS. These results suggest that upon stimulation these animals are more prone to UCP1 activation and potentially able to increase thermogenesis. Future work should address the effect of altered mitochondrial dynamics in the NTS on the thermogenic activity of the BAT.

We observed substantial lipid droplet enlargement in the BAT of GFAP-GFP control animals under HFD diet, and this was associated with a significant higher level of *Cidea* mRNA transcripts compared to GFAP-Drp1-K38A animals. This gene is associated with lipid droplet enlargement and is essential for adipocyte differentiation and regulation of lipolysis and lipogenesis coupling in BAT [48]. The lower levels of *Cidea* we observed in GFAP-Drp1-K38A animals could play a role in the prevention of BAT lipid droplet enlargement. *Cidea* transcripts, however, were also significantly lower in GFAP-Drp1-S637A animals compared to matching GFAP-RC controls; whilst no effects on BAT morphology were seen in absence of HFD. It would therefore be of interest to investigate the significance of this finding in the context of activation of mitochondrial fission in astrocytes of the NTS in GFAP-Drp1-K38A-expressing animals fed with RC.

There are direct connections between the NTS and the dorsal motor nucleus of the vagus (DMX) where vagal efferent are stimulated to send signals to the peripheral organs, a potential direct modulation of the BAT via vagal efferent could be also possible. Further investigation, maybe with retrograde tracing, would be required to determine how BAT activity is modulated by the NTS.

Data in this study were obtained from male rats. It is well documented that males and females respond in different ways to hormonal stimulation and diet, for example, while in male rats, insulin decreases food intake, in females it does not [49–51]. This was also shown in humans treated with intranasal insulin [52]. Future work should address the effect that HFD-feeding and altered mitochondrial dynamics in the NTS have on female rats.

In humans, individuals with preserved BAT have a lower prevalence of cardiometabolic diseases, and lower odds of developing type 2 diabetes, dyslipidemia, coronary artery disease, cerebrovascular disease, congestive heart failure and hypertension. They also have better blood glucose and cholesterol regulation. Interestingly overweight and obese individuals with detectable BAT presented the aforementioned beneficial effects thus suggesting that possessing healthy BAT is important to prevent the deleterious effects of obesity and potentially protect from developing diabetes [53]. Our data show that alteration of BAT activity and accumulation of WAT-like fat droplets occurs within two weeks of HFD-feeding and that the NTS plays a fundamental role in preserving BAT function in this HFD-fed model. Therefore, our research indicates that the NTS might be a novel target for the treatment and prevention of obesity and diabetes and for preserving general cardiometabolic health.

## 4. Methods

### 4.1. Animal models

Nine-week-old wild-type Sprague Dawley male rats weighing 245±15g were obtained from Charles River Laboratories (Margate, UK) and were used in line with the United Kingdom Animals (Scientific Procedures) Act 1986 and ethical standards set by the University of Leeds Ethical Review Committee. Animals were individually housed and maintained on 12-hour light-dark cycle from 6 AM to 6 PM with access to standard regular chow (RC), high-fat diet (HFD) or control (RC) diet *ad libitum*, with the exclusion of the pair-fed cohort. On day 0 rats were stereotactically implanted with a bilateral cannula targeting the NTS within the DVC (0.0 mm on the occipital crest, 0.4 mm lateral to the midline and 7.9 mm below the skull surface). On day 1 an adenoviral system was used to deliver either a flag-tagged dominant-negative form of Drp1 (CMV-Drp1-K38A or GFAP-Drp1-K38A), a Flag-tagged constitutively active form of Drp1 (CMV-Drp1-S637A or GFAP-Drp1-S637A) or a control green fluorescent protein (CMV-GFP or GFAP-GFP) under the control of the cytomegalovirus (CMV)[22] or GFAP[23] promoters. Control groups expressing CMV-GFP or GFAP-GFP were pulled together since the effects were comparable, this allowed us to decrease the number of animals used according to the NC3R.

The diet compositions were as follows: control RC (3.93 kcal/g) was 61.6% carbohydrate, 20.5% proteins, 7.2% fat and 3.5% ash (F4031), HFD (5.51 kcal/g) was 36.2% carbohydrate, 20.5% protein, 36% fat, 3.5% ash (F3282) both Datesand ltd. (Manchester, UK). Animals were either submitted for PET/CT scans and sacrificed or directly sacrificed on day 16 and brown adipose tissue (BAT) and epididymal, retroperitoneal and visceral fat were collected and weighed. Liver, plasma, and BAT were collected and stored for further analysis. DVCs were collected for immunohistochemistry to confirm successful targeting in surgery. Rats that did not recover after surgery (lost more than 10% of their body weight) were not used in the experiments. Rats that at the end of the experiment did not show the viral expression in the NTS were removed from the analysis.

### 4.2. Generation of adenoviral vectors

Adenoviruses expressing FLAG-Drp1K38A, FLAG-Drp1-S637A and GFP under the CMV promoter were generated as described in Filippi et al. (2017)[22]. To produce an adenoviral system expressing the mutant proteins under the GFAP promoter, the CMV promoter was removed from the pacAD5 shuttle vector, and replaced with the rat GFAP promoter. Recombinant adenoviruses were amplified in HEK 293 AD cells and purification was performed using a sucrose-based method [54]. Adenoviruses injected into the animals’ brains had a titre between 3 x 109 pfu/ml and 4.4 x 1011 pfu/ml.

### 4.3. Pair-fed studies

The GFAP-Drp1-K38A animals were used as control animals and given ad libitum food at all times. Pair-fed GFAP-GFP rats were given the average amount of food the control animals had consumed on the previous day. The cumulative weight gain and food intake of the animals were monitored daily.

### 4.4. PET/CT scans

Scans were performed at the Experimental and Preclinical Imaging Centre (ePIC) at the Leeds Institute of Cardiovascular and Metabolic Medicine (LICAMM) and 2-deoxy-2-^18^F-fluoro-β-D-glucose (^18^F-FDG) was synthesised at Royal Preston Hospital (Preston, UK) and delivered on the day of the scans. 30 minutes prior to scans, animals were injected intraperitoneally with 1mg/kg of adrenergic agonist CL 316,243 in 0.9% saline solution or 0.9% saline vehicle. Animals were then induced to a surgical plane of anaesthesia with 5% isoflurane at 2L/min and a bespoke line was inserted in the tail lateral vein for ^18^F-FDG administration. ^18^F-FDG was prepared in 500 µl of 0.9% saline solution (AquaPharm, Animalcare Ltd, York, UK) and 21.86±0.73 (n=19) Mbq was injected simultaneously to the initiation of the dynamic scan. PET/CT scans were acquired on an Albira Si PET-CT system (Bruker, Billerica, MA, USA) using a combined 90-minute single PET and a CT Triple Standard (Low Dose, Low Voltage) sequence. The animals were placed prone in the scan bed and the centre of view was positioned on the BAT. Animals were kept at 2.5% isoflurane during the duration of the scan. At the end of the scan (t=4800 s) animals were sacrificed by decapitation and tissues harvested for gamma counting. ^18^F-FDG tissue absorption in the gamma counting assay was calculated as percentage of the total dose of injected ^18^F-FDG per gram of tissue corrected by ^18^F decay (half-life t=109.7 s), which included correction for residual activity in the syringe following injection, and IV catheter prior to sacrifice. After the PET scan, CT images were obtained using the following parameters: x-ray energy 40 kVp, resolution 90 µm, 360 projections, 8 shots. CT data was acquired using the three-dimensional ordered subset expectation maximisation algorithm.

PET and CT scans were reconstructed using the Albira software (Bruker, Billerica, MA, USA). CT was reconstructed using the Filtered Back Projection (FBP) algorithm set on high. The PET images were generated using the Maximum Likelihood Estimation Method (MLEM) set at 0.25 mm and partial volume correction and attenuation and activation of scatter randoms and decay corrections. Reconstructed images were fused and visualised using the pmod software (pmod technologies, Zurich, Switzerland) with SUV set at 3. PET scans were aligned to CT, filtered with a Gaussian filter (1mm) and analysed by comparing three-dimensional regions of interest (VOI) of brown adipose tissue to the inferior lobe of the right lung. Time-activity curves for the ROI were generated using the VOI statistics function of the pmod software.

CT scans were analysed to quantify total visceral and subcutaneous volumes. The fat segmentation method [55] was used by employing Hounsfield units (HU). Total body volume was segmented at –300 HU, +3500 HU and adipose tissue was segmented at – 190 HU, -60 HU and the total percentage of fat volume mass was calculated. Animals that did not express the viruses in the NTS were excluded from the analysis.

### 4.5. Hematoxylin and Eosin

BAT pads were dissected and post-fixed in 4% PFA. Paraffin embedding, cutting and H&E staining steps were performed by the Division of Pathology at St James University Hospital (Leeds University Teaching Hospitals, UK). Sections were examined and imaged by light microscopy at 40x magnification using an Evos imaging system (Invitrogen, Waltham, MA, USA) and analysed using Image J (National Institute of Health, Bethesda, MD, USA). The total number and size of lipid droplets per field of view was obtained by assuming droplets’ circularity to calculate their area. Area cut-off points for brown adipocytes were calculated from their maximum acceptable size according to the literature of 20 µm [56]. Images were then converted to 16-bit and segmented by threshold filtration at a fixed value and made binary using the water-shed function to reconstruct lost or lacerated cellular membranes. Lastly, the particle analysis tool was used to obtain average adipocyte size and adipocyte numbers. Adipocytes bordering the ROI frame were excluded.

### 4.6 Immunohistochemistry

BAT pads were dissected and post-fixed in 4% PFA for 2 hours, and animals were perfused with 4% PFA to fix the brain. The brainstem areas containing the DVC and BAT sections were put in 15% followed by 30% sucrose until saturated and frozen in a cryo-embedding medium (Leica Biosystems, Nussloch, Germany). Brain sections were cut at 30 µm and BAT sections were cut at 5 µm using a cryostat (Leica Biosystems, Nussloch, Germany). Brain sections were labelled with Anti-FLAG M2 F1804) (1:500) (Sigma, Burlington, MA, USA), and co-stained with the neuronal marker NeuN (ABN90) (1:2000) (Millipore, Burlington, MA, USA) and astrocytic marker GFAP (ab7260) (1:1000) (Abcam, Cambridge, UK). BAT was stained with an antibody against tyrosine hydroxylase (TH) (ab113) (1:250) (Abcam, Cambridge, UK). Host specific AlexaFluor secondary antibodies were used at 1:1000. BAT images for analysis were acquired using an Evos imaging system (Invitrogen, Waltham, MA, USA) and representative images for brain and BAT were taken using a Zeiss LSM800 upright confocal microscope (ZEISS, Wetzlar, Germany). 4-5 images from n=4 animals per group were selected for TH quantification. This was done using a threshold function, and further confirmed with particle analysis to isolate TH pools-which are defined as the accumulation of TH-within nerve terminals within BAT. Both analyses were carried out using Image J (National Institute of Health, Bethesda, MD, USA).

### 4.7. RNA extraction and cDNA synthesis

After sacrifice, samples were snap frozen and weighed and RNA was extracted from 30∼50 mg of BAT. Samples were homogenised in 1 ml QIAzol lysis reagent (Qiagen, Hilden, Germany) with a handheld tissue homogeniser (Appleton Woods Ltd, Birmingham, UK). RNA was extracted using the RNeasy Lipid Tissue Mini Kit (Qiagen, Hilden, Germany) as per the manufacturer’s instructions, including a DNAse digest step. RNA integrity analysis was performed using an RNA 6000 Nano Kit and run on a 2100 Bioanalyser, (Agilent, Santa Clara, CA, USA). Samples with a 260/230 score of 2-2.2, 260/280 score of 1.8-2.2 and a RIN score of at least 8 were used for cDNA synthesis and further downstream analysis. Single-stranded cDNA was synthesised from 100 ng of RNA and messenger RNA was selectively amplified in the first step of the reaction using Oligo-(dT)20 and dNTPs (both ThermoFisher Scientific, Waltham, MA, US). In the second reaction step, Superscript III kit (ThermoFisher Scientific, Waltham, MA, US) and RNAsin Plus (Promega, Southampton, UK) were used to carry out cDNA synthesis according to manufacturer’s instructions.

### 4.8. Quantitative real-time PCR

All samples were run in triplicates. RT-qPCR reactions were carried out using a SYBR green master mix (Applied Biosystems, Waltham, MA, USA) on a QuantStudio 3 qPCR system (ThermoFisher Scientific, Waltham, MA, US).*36b4* gene encoding for acidic ribosomal phosphoprotein P0 was selected as house-keeping control due to its high stability in adipose tissues (Zhang et al., 2016). qPCR reactions containing primers for genes of interest and a serial cDNA dilution of sample pool (1:50, 1:100, 1:200) were performed using combined comparative Ct and melting curves to ensure primers quality and test optimal cDNA dilution. Following this, the qPCR products were run on a 4% agarose gel and analysed to ensure cDNA product integrity, confirm the predicted *in silico* size for the primers and screen for potential primer-dimer formation (Supplementary figure 5).

### 4.9. Enzyme-linked Immunosorbent assay (ELISA) for plasma insulin

At the end of the experimental work, animals were sacrificed by pentobarbital overdose (60 mg/kg) and whole blood was collected and spun down to separate the plasma and obtained samples were snap frozen until analysis. A rat insulin ELISA kit low range assay (0.1-6.4 ng/ml) (Crystal Chem, Elk Grove, IL, USA) was used to measure insulin plasma levels according to manufacturer instructions.

### 4.10. Coenzyme Q (CoQ) Competitive ELISA

To measure Coenzyme Q (CoQ) levels in BAT a commercial Rat Coenzyme Q ELISA KIT (MBS7241152) (My BioSource, San Diego, CA, USA) was used according to manufacturer instructions.

### 4.11. Statistical analysis

All data are expressed as mean ± SEM, and where applicable data from each animal is reported as a single data point. All collected data was tested for normality prior to analysis using the Shapiro-Wilk normality test, and analysed using Prism 9 Software (GraphPad, San Diego, CA, US). Significant differences were determined by unpaired t-test, one-way ANOVA (Post-hoc test: Tukey), or a two-way ANOVA (Post-hoc test: Tuckey). T-test was used for tissue mass, histology and plasma analysis and qPCR; One-way ANOVA for PET/CT scan data and Two-way ANOVA for chronic feeding studies. The prism Outlier calculator was used to identify and appropriately exclude outliers, if present. In Figure legends and text, n refers to the number of animals used in each experiment, and when appropriate the number of technical replicates is also reported. P < 0.05 was statistically significant. Significance was defined by (∗) P < 0.05; (∗∗) P < 0.01; (∗∗∗) P < 0.001; (∗∗∗∗) P < 0.0001. where indicated other symbols like $, &, ^ and # were used together with the *.

## Acknowledgment

This work was supported by grants from the MRC, MRC-Career Development Fellowship (MR/S007288/1) and the Medical Research Foundation (MRF, MRF-167-0001-RG-FILI-C0838). Microscopy was performed in the bioimaging facility in Leeds and the Zeiss LSM 880 inverted confocal was funded by Wellcome Trust (104918/Z/14/Z). PET scanning was performed in the Experimental and Preclinical Imaging Centre (ePIC) funded by the BHF (SI/14/1/30718). We acknowledge technical support from Joanna Koch-Paszkowski and Simon Futers for conducting the PET scans. H&E staining was performed by the Histology facility at the Wellcome Trust Brenner Building St. James’s University Hospital Leeds. Graphical abstract was created with BioRender.com.

## Author contributions

**A.F.** conducted and designed experiments, performed data analyses and wrote the first draft of the manuscript. **L.E. N**. Conducted in vivo experiments. **J.C.G.** Assisted with the experiments. **B.P.** Assisted with experiments. **S. A. D.** Contributed in experimental design, and discussion of results and critiqued the manuscript. **B.M.F.** conceived and supervised the project, designed and conducted the experiments, and wrote the manuscript.

## Supplementary Material

**Supplementary Figure S1:**
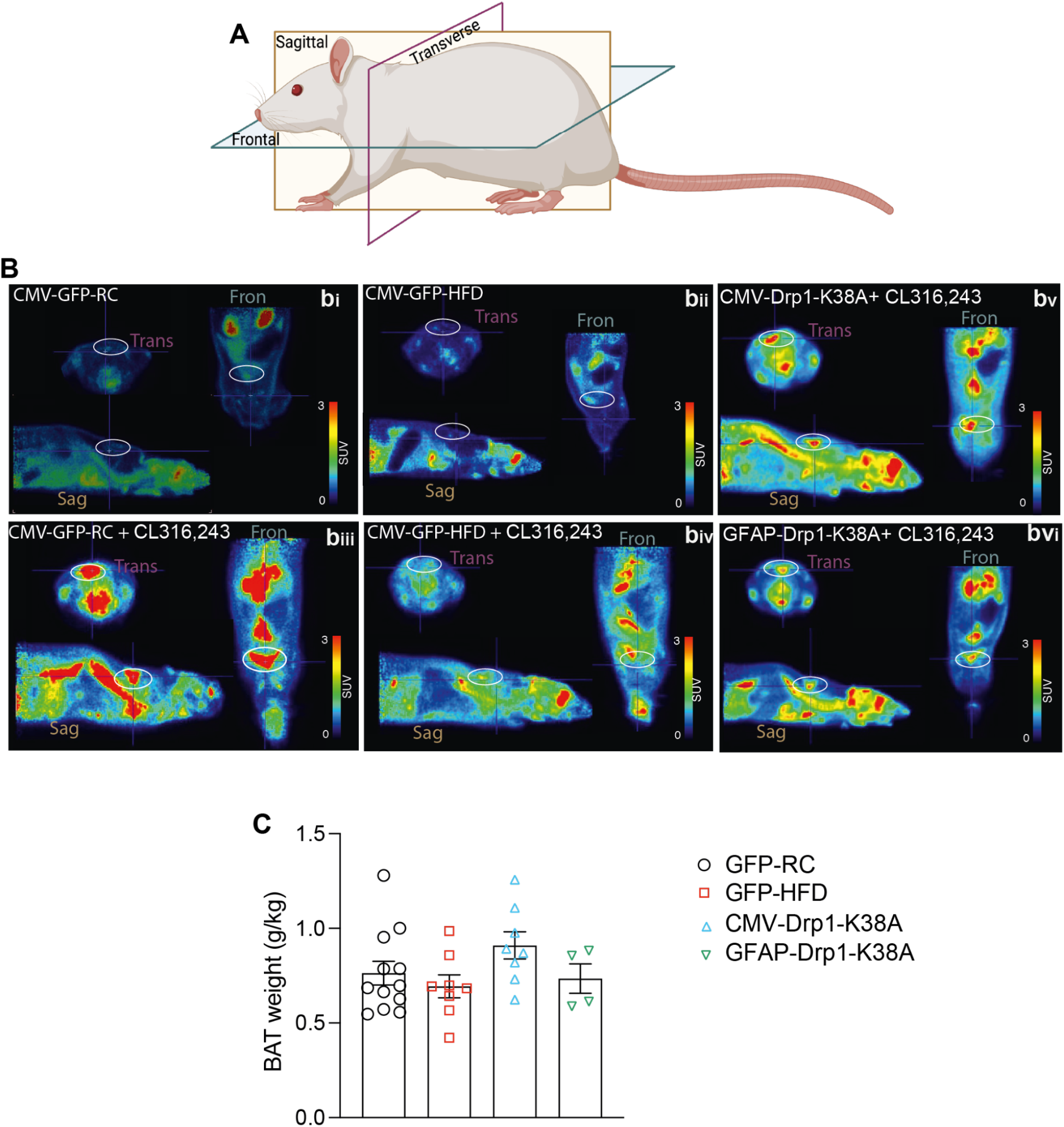
Representative PET scan images. **(A)** Schematic showing the different anatomical planes in the rat. **(B)** representative PET images of the different PET scans (Names in the panel) showing 3 different anatomical planes with BAT area circled. Panel **bi to biv** refers to Figure 1E to H; panel **biv to bvi** refers to Figure 1 K to M**. (C)** BAT weight normalised to body weight (g/kg) in GFP expressing RC (n=11) or HFD (n=8)-fed rats and in HFD-fed rats expressing CMV-Drp1-K38A (n=8) or GFAP-Drp1-K38A (n=5) BAT weight is taken from rats that received PET scan in Figure 1. Abbreviations Trans - transverse, Fron - frontal, Sag - Sagittal

**Supplementary Figure S2.**
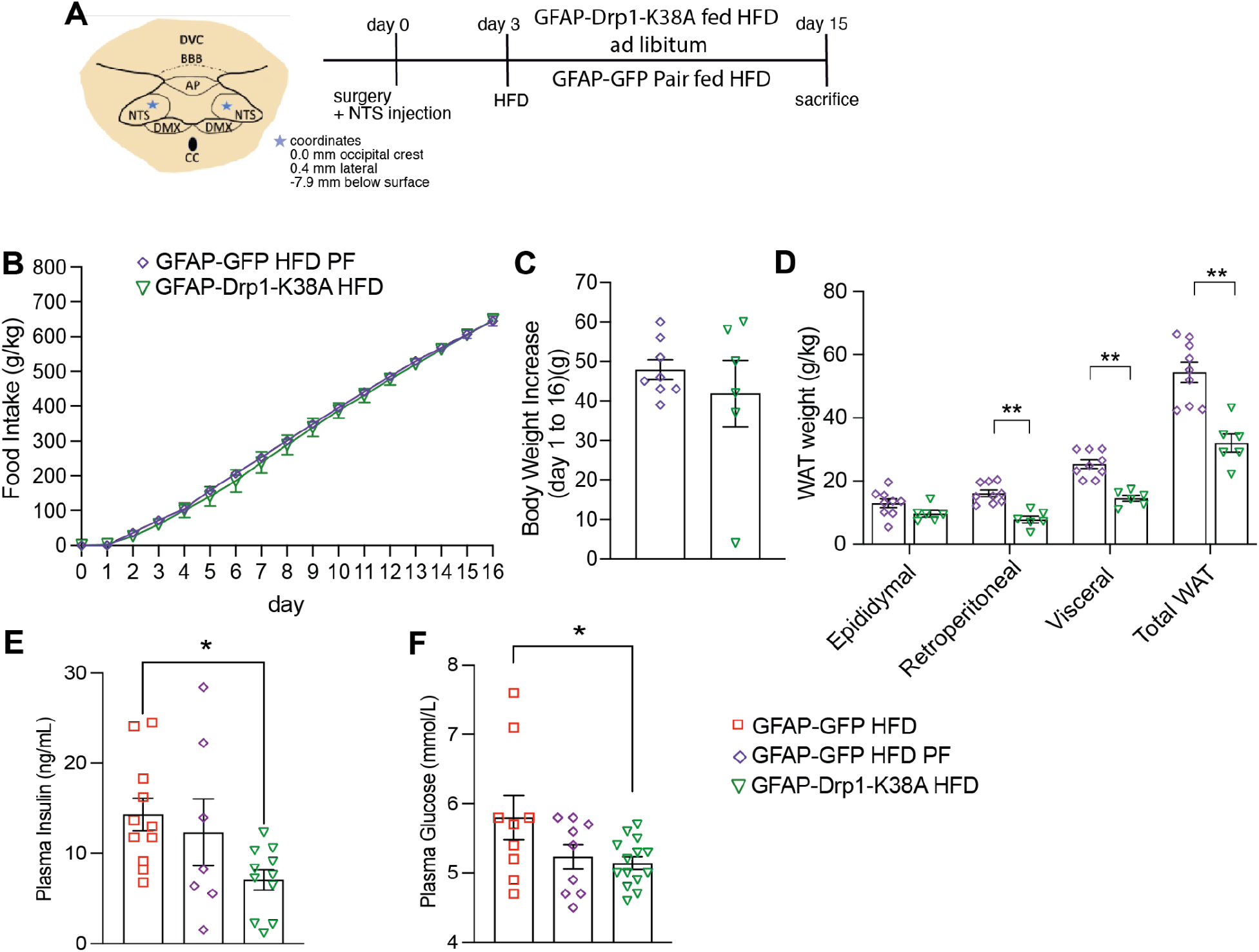
Food intake, body weight, WAT, plasma insulin and blood glucose levels from the HFD ad-libitum and pair-fed cohorts. (A) The experimental design of the 2 weeks feeding study. (B) Food intake of GFAP-GFP HFD PF (n=8) and GFAP-Drp1-K38A HFD (n=6) normalised to body weight in kg. (C) Body weight increase of GFAP-GFP HFD PF (n=8) and GFAP-Drp1-K38A HFD (n=6) at day 16 of the feeding study. (D) WAT weight normalised to body weight (kg) for each abdominal white adipose tissue depot. (E) Plasma insulin levels for GFAP-GFP HFD (n=11), GFAP-GFP HFD PF (n=7) and GFAP-Drp1-K38A HFD (n=11). (F) Blood glucose for GFAP-GFP HFD (n=9), GFAP-GFP HFD PF (n=8) and GFAP-Drp1-K38A HFD (n=14). All data was tested for normality prior statistical tests using the Shapiro-Wilk normality test. Statistical test: Two-way ANOVA, post-hoc Sidak (B), t-test (C-D) or One-way ANOVA, post-hoc Sidak (E-F). ∗p < 0.05; ∗∗p < 0.01; ∗∗∗p < 0.001; ∗∗∗∗p < 0.0001. Values are shown as mean ± SEM and single data points are highlighted.

**Supplementary Figure S3.**
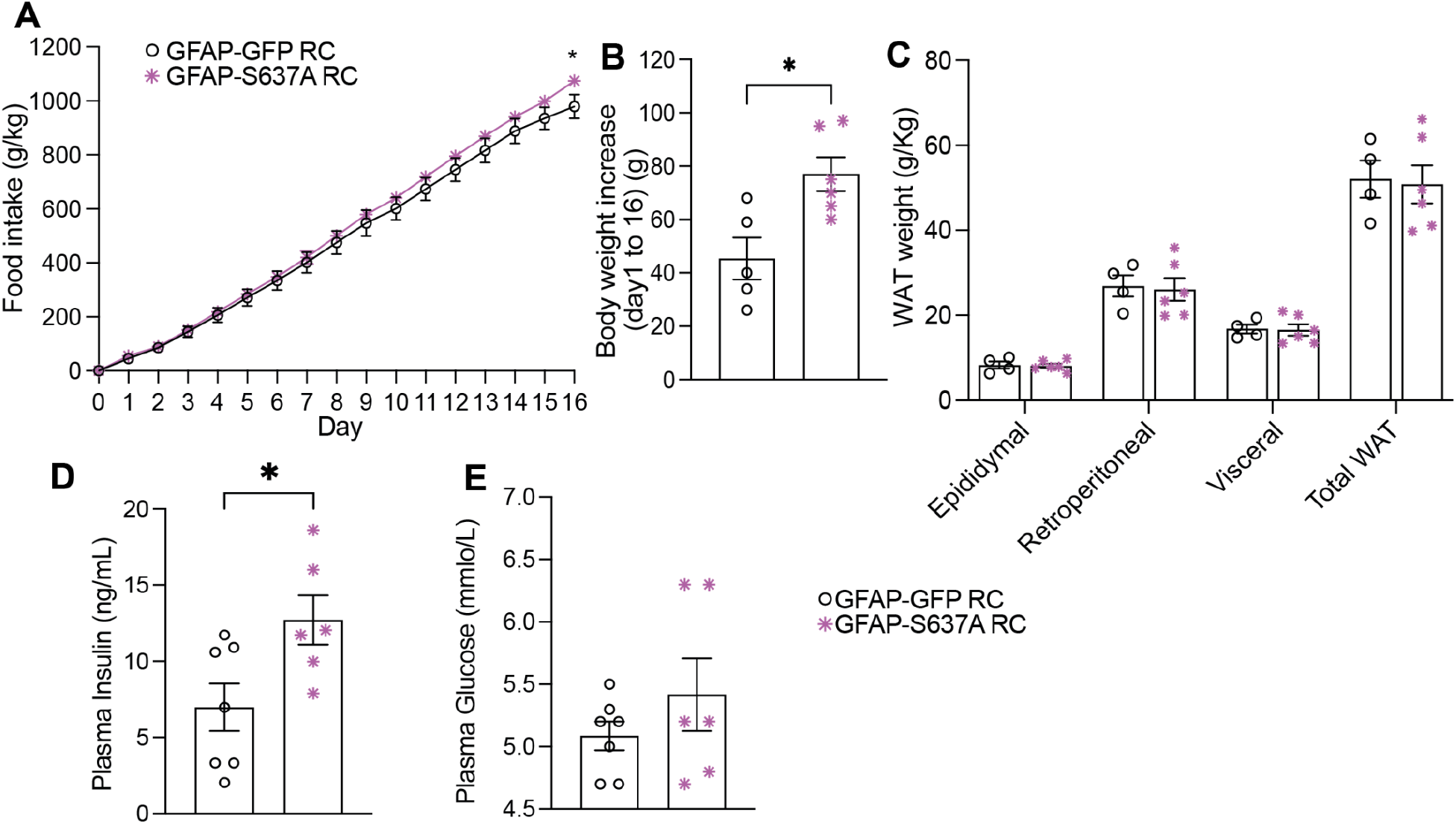
Food intake, body weight, WAT, plasma insulin and blood glucose levels from the GFAP-Drp1-S637A RC cohort. (A) Food intake of GFAP-Drp1-S637A RC (n=6) and GFAP-GFP RC (n=5) normalised to body weight in kg. (B) Body weight increase of GFAP-Drp1-S637A RC (n=6) and GFAP-GFP RC (n=5) animals at day 16 of the feeding study. (C) WAT weight normalised to body weight (kg) for each abdominal white adipose tissue depot. (D) Plasma insulin levels for GFAP-Drp1-S637A RC (n=6) and GFAP-GFP RC (n=7). (E) Blood glucose level for GFAP-Drp1-S637A RC (n=6) and GFAP-GFP RC (n=7). All data was tested for normality prior statistical tests using the Shapiro-Wilk normality test. Statistical test: Two-way ANOVA, post-hoc Sidak (A), t-test (B-E). ∗p < 0.05; ∗∗p < 0.01; ∗∗∗p < 0.001; ∗∗∗∗p < 0.0001. Values are shown as mean ± SEM and single data points are highlighted.

**Supplementary Figure S4.**
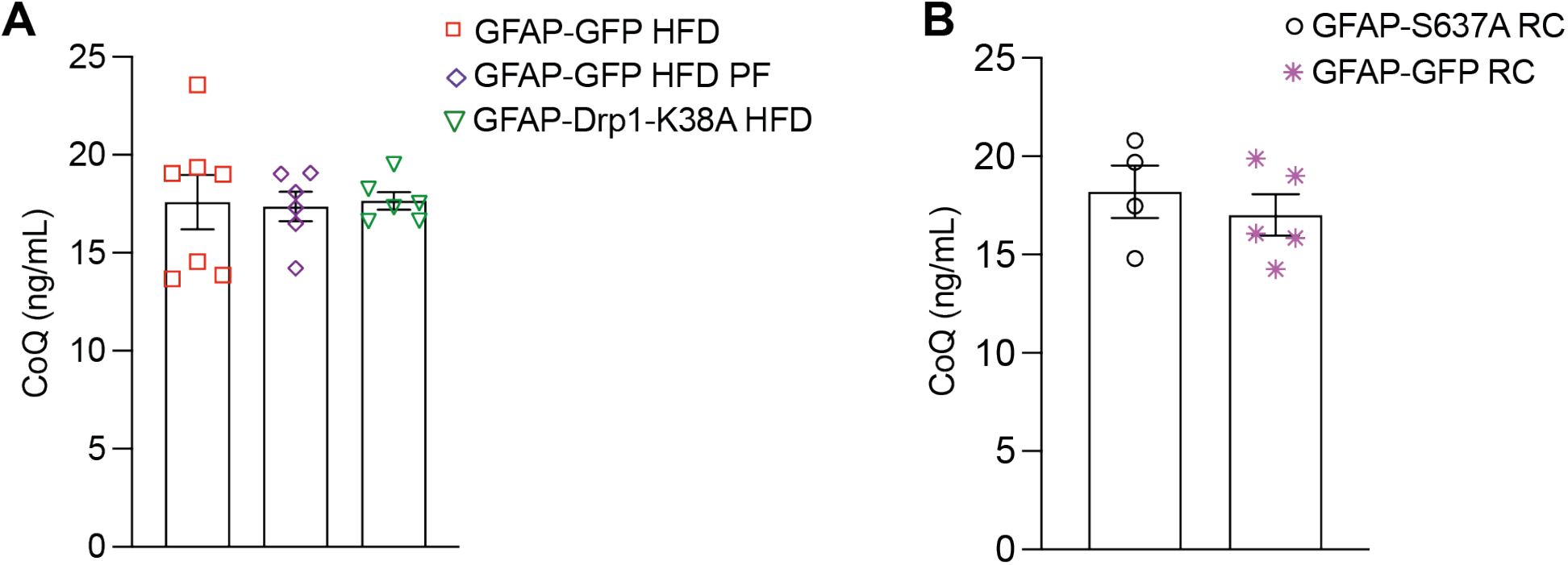
CoQ level measured in all cohorts. (A) CoQ in ng/mL measured in GFAP-GFP HFD (n=7), GFAP-GFP HFD PF (n=6) and GFAP-Drp1-K38A (n=7). (B) CoQ in ng/mL measured in GFAP-GFP RC (n=4) and GFAP-Drp1-S637A (n=5). All data was tested for normality prior statistical tests using the Shapiro-Wilk normality test. Statistical test: One-way ANOVA, post-hoc Tukey(A) or T-Test (B). Values are shown as mean ± SEM

**Supplementary Figure S5:**
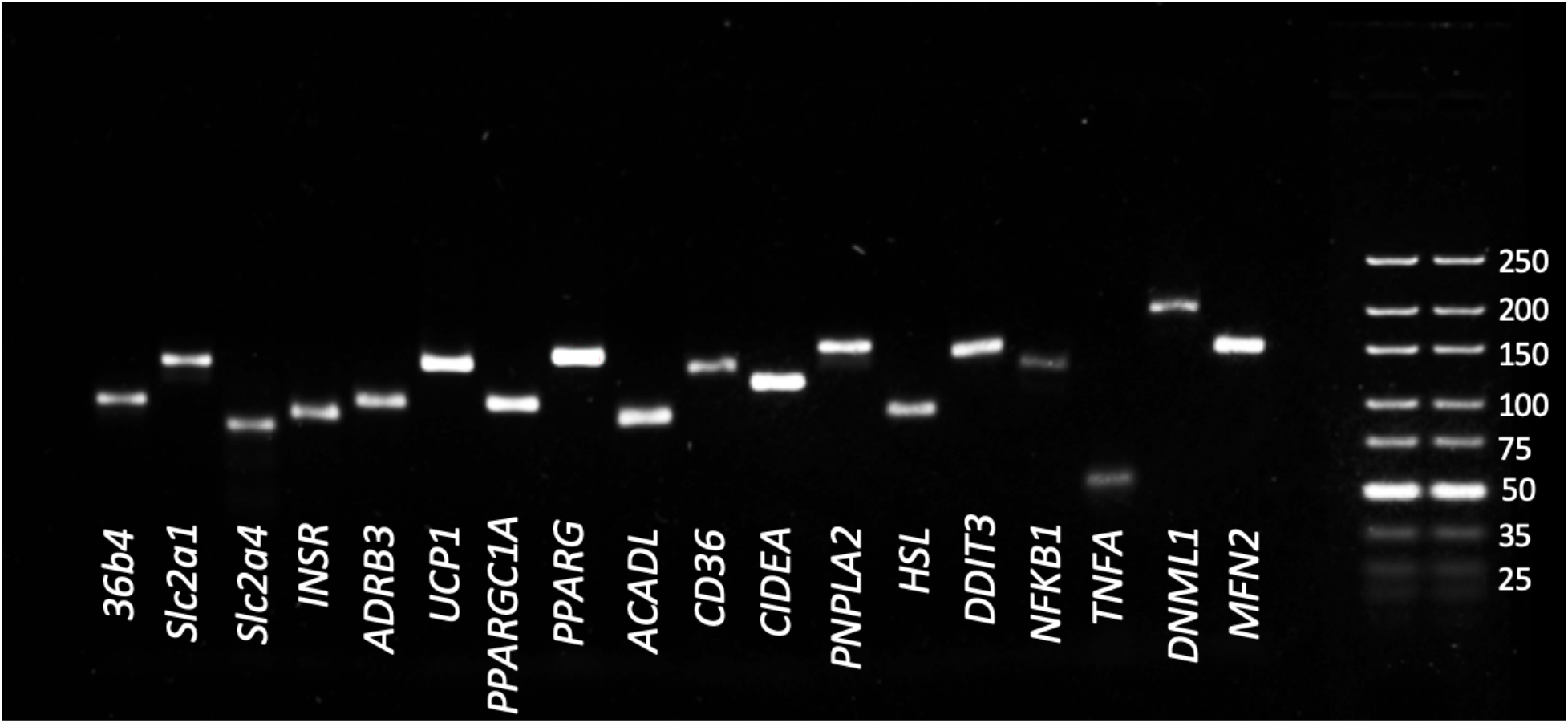
Representative agarose gel showing that the size of the primers employed in this study correspond to the *in silico* predicted size and that they produce a single clean product free of smears or primer-dimer formations.

